# Limited codiversification of the gut microbiota with humans

**DOI:** 10.1101/2022.10.27.514143

**Authors:** Benjamin H. Good

## Abstract

Gut bacteria exhibit striking variation across different human populations, but the evolutionary forces that have shaped this diversity are less well understood. Recent work has argued that many species of gut bacteria have codiversified with modern humans, based on the phylogenetic correlations between human and microbial genomes. Here we reanalyze these data and show that the correlations between human and microbial phylogenies are often substantially weaker than between unlinked human chromosomes, and that similar correlations can arise through geographic structure alone. These results suggest that traditional codiversification has been limited in recent human history, and highlight alternative strategies for quantifying the extent of human-microbe coevolution.

## Main Text

The human microbiome is highly diverse, with individuals varying in both the species they contain (1) as well as the genetic variants (or “strains”) of each species (2, 3). Advances in metagenomic sequencing have made it possible to catalog this intra-species variation in human populations from around the world (4–9). Understanding the origins and implications of this diversity – and how it depends on the lifestyle and ancestry of the host – is an important frontier of microbiome research.

A recent study by Suzuki & Fitzstevens *et al* (9) shed new light on this problem by assembling a large collection of sequenced human gut metagenomes and paired host genomes from several industrialized populations around the world. The authors used these data to argue that dozens of species of gut bacteria have evolved in parallel (or “codiversified”) with modern humans, based on the observed genetic correlations between human and microbial genomes (9). An accompanying perspective (10) proposed a stronger model, in which gut bacteria may have kept fidelity to human lineages for thousands of human generations. Such intimate commensal relationships would be remarkable if true, since they would create extensive opportunities for coevolution between human and bacterial genomes.

While striking examples of codiversification have been observed between host species (11, 12), they are more challenging to detect on the shorter evolutionary timescales separating modern human populations. The underlying genetic distance scales are roughly compatible: estimates of the mutation accumulation rate in human gut bacteria (13, 14) suggest that isolated bacterial lineages should have diverged by ∼1-10% in the ∼60,000 years since the out-of-Africa expansion (15). These divergence levels are comparable to the observed synonymous diversity within many species of human gut bacteria (16), potentially hinting at a common evolutionary origin.

However, similar to our own genomes, the genetic diversity of the gut microbiota is highly intermingled across space and time. Previous work has shown that the divergence between strains from different continents is often comparable to the diversity among strains from the same geographic location (16, 17). Horizontal transmission is also widespread. Unrelated hosts from different continents can sometimes share nearly identical bacterial strains (18–20), while even the most closely related hosts – identical twins – are often colonized by highly diverged strains over their lifetimes (19).

To identify signatures of codiversification against this backdrop, Suzuki & Fitzstevens *et al* (9) compared genome-wide phylogenies constructed from human and microbial genomes. Care must be taken when interpreting such trees on shorter within-species timescales: recombination within both humans and gut bacteria (19, 20) produces mosaic patterns of ancestry that are rarely captured by a single phylogenetic tree. However, recent work has shown that the genome-wide phylogenies inferred from recombining populations can still encode some information about the historical patterns of gene flow within a species (21). In these settings, similarities between the human and bacterial phylogenies will reflect the migration patterns of larger ancestral populations (17), rather than the genealogical fidelity of individual microbial lineages.

Suzuki & Fitzstevens *et al* (9) used a phylogenetic congruence test (PACo) to quantify the similarities between the human and bacterial phylogenies. By permuting the links between hosts and bacteria, they found that 36 of 59 bacterial species were more similar to the human phylogeny than expected by chance (q<0.05). While these significant q-values show that the human and bacterial phylogenies are not completely independent, they do not directly demonstrate the evolutionary parallelism implied by traditional models of codiversification (22). For example, theoretical arguments show that small amounts of geographic structure can also produce significant PACo scores even in the absence of codiversification (Fig. 1), since human genetic diversity is already correlated with geography (23). The presence of these confounding factors suggests that additional information is needed to interpret the PACo scores observed in Ref. (9).

**Figure 1:**
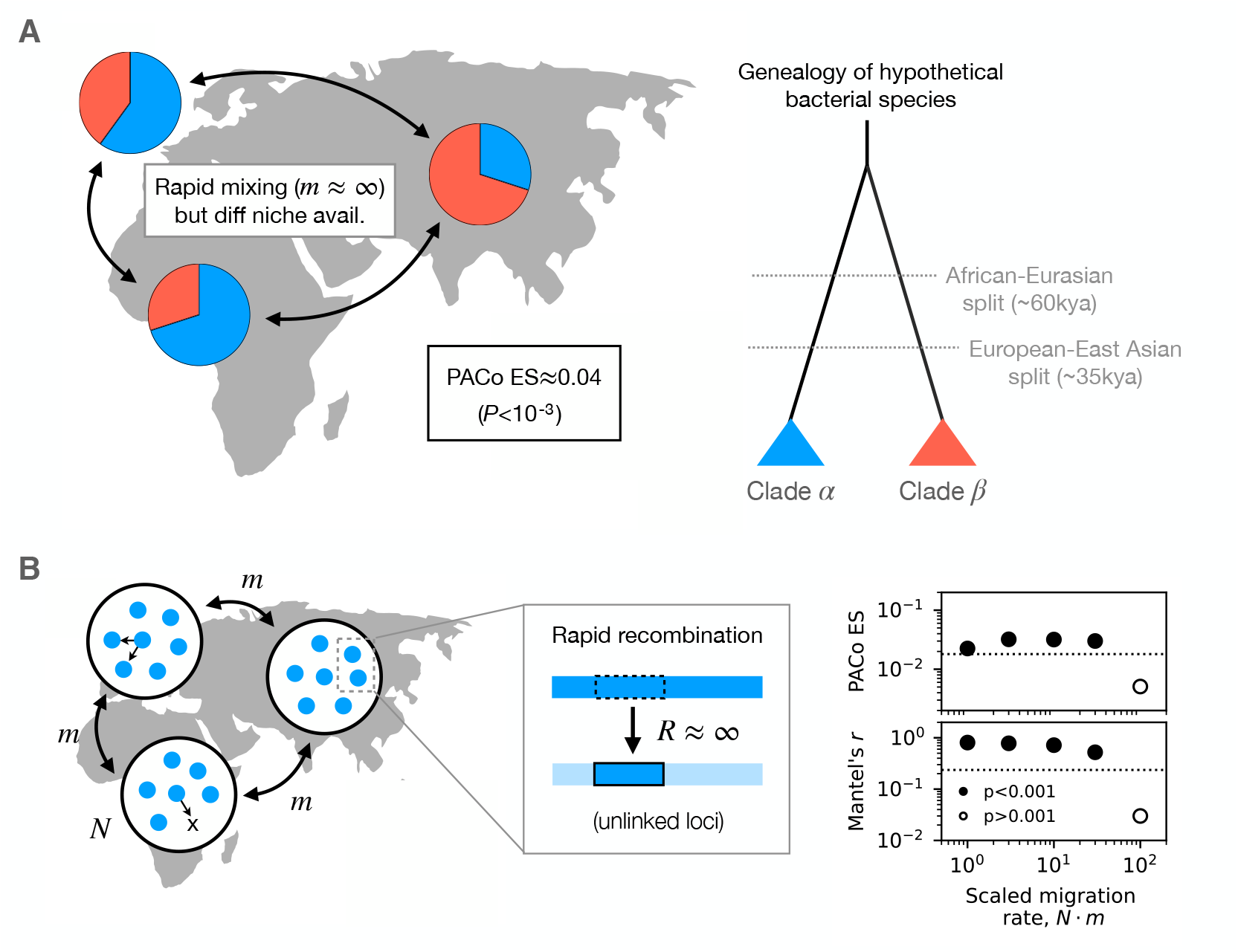
Geographic structure can produce statistically significant phylogenetic correlations even in the absence of codiversification. (a) An example of a hypothetical bacterial species that did not evolve in parallel with modern humans. Two genetically isolated clades originated before the out-of-Africa migration (right), and are currently shared among all human populations. Small shifts in the frequencies of the clades across continents (unrelated to human ancestry) can produce spurious correlations with the observed human phylogeny (left) that are comparable to the largest effect sizes observed in Ref. (9) (Fig. 2D); PACo scores were calculated using the observed human phylogeny for the subset of hosts in Ref. (9) that contained strains of *Prevotella copri*. (b) An alternative spatial null model with finite rates of migration and widespread within-species recombination. Recombination occurs infinitely quickly within each continent, leading to rapid unlinking of mutations. Genome-wide phylogenies inferred from simulations of this model exhibit significant correlations with the observed human phylogeny (Methods), even when most strains recently migrated from another continent (*mT*_mrca_ ≳ *Nm* ≫ 1). Points denote simulation results and their associated p-values, while the dashed lines show the corresponding levels observed for *P. copri* (Methods). These theoretical examples show that phylogenetic correlations do not necessarily imply codiversification.

In principle, independent regions of the human genome could provide a positive control for genomes that have truly codiversified with each other. While mitochondria are strictly maternally inherited, different halves of the nuclear genome trace their ancestry back to hundreds of distinct genetic ancestors within a few hundred years, similar to a large random sample from the same local population (24). The residual correlations between these chromosomes can therefore provide a natural null model for codiversification that does not rely on strict maternal inheritance. However, the same PACo analysis performed on separate halves of the human genome yields an effect size of ES≈ .7 – which is substantially greater than the largest effect sizes reported in Ref. (9) (Fig. 2D). Similar values are obtained when the host labels are randomized within countries (ES≈0.6) or continents (ES≈0.54), confirming that they are driven by the largest geographic scales. This large gap in effect sizes suggests that the parallelism observed between human and bacterial phylogenies is substantially lower than expected under traditional models of codiversification.

**Figure 2:**
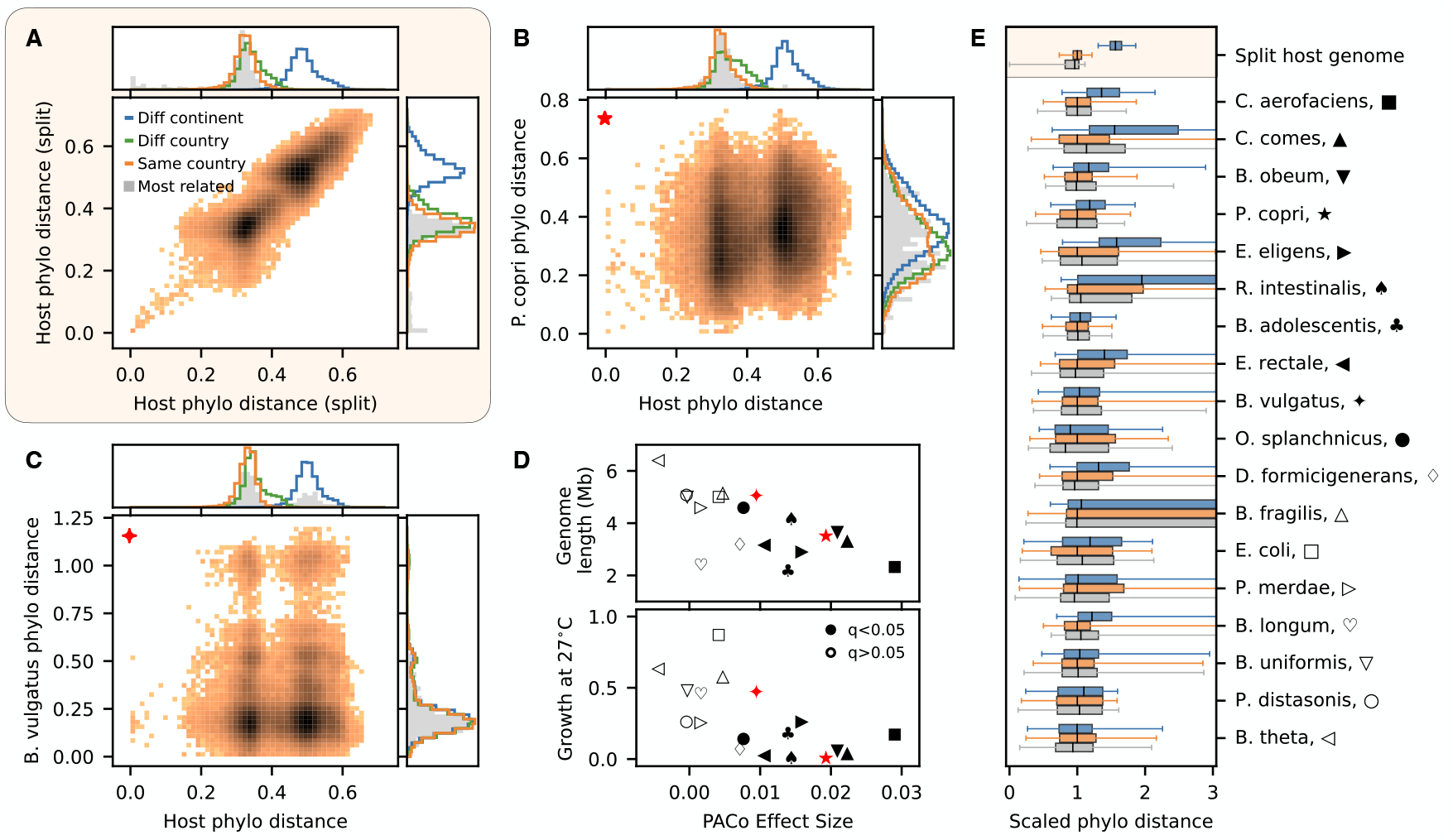
Host genetic similarity is not strongly predictive of bacterial genetic similarity. (A-C) Phylogenetic correlations for separate halves of the human genome (A), *Prevotella copri* (B), and *Bacteroides vulgatus* (C). Heat map shows estimated phylogenetic distances between pairs of adult individuals sequenced in Ref. (9); three samples with anomalously large host distances (Fig. S2) were removed so that they would not bias the correlation in panel A. Right histogram panel shows the distribution of bacterial distances for hosts in different geographic categories (blue, orange, green), or in the next most closely related hosts (grey); top histogram panel shows analogous distributions for host phylogenetic distance, using the next most closely related bacterial strain to construct the grey distribution. (D) Correlation between PACo effect size and genome length (top) or relative growth at 27^*°*^C (bottom) as measured by Ref. (9). *P. copri* and *B. vulgatus* are indicated in red. (E) Summarized versions of the right histograms in panels A-C for all of the species in panel D. Box plots show medians and interquartile ranges (boxes) and 95% confidence intervals (whiskers); phylogenetic distances are rescaled by the median within-country distance for each species.

These results are not specific to the PACo test. Direct examination of the phylogenetic distances reveals that two halves of the human genome are substantially more correlated with each other than with any gut bacterial genomes (Fig. 2). For species with the strongest reported codiversification scores (e.g. *Prevotella copri*, Fig. 2B), one can observe a small but significant decrease in bacterial relatedness with increasing host distance (P<10^-3^; Mantel test). However, these small average differences are dwarfed by the strain variation within individual countries (Fig. 2B, right, orange) or among the most closely related hosts (grey). Similar behavior can be observed in other species with significant PACo scores (e.g. *Bacteroides vulgatus*, Fig. 2C), though the subtle shifts in mean relatedness can be more difficult to discern by eye (P<10^-3^, Kolmogorov-Smirnov test). In each of these cases, even the most closely related hosts contain bacteria that span nearly the full range of bacterial relatedness, while hosts from different continents frequently share strains that are as similar as those from the most closely related hosts (Fig. 2E, Figs. S1 & S2). These findings are consistent with previous observations in adult twins (19), suggesting that they do not sensitively depend on the phylogenetic inference scheme employed here (Figs. S3 & S4).

Together, these results suggest that traditional codiversification has been limited within modern human populations. Gut bacteria may have still adapted to specific groups or geographic locales. But for most of the species and populations examined here, the genome-wide diversity of their constituent strains does not strongly mirror the evolutionary history of their host populations.

This distinction has important evolutionary implications, and also potential practical consequences. Strong forms of codiversification might imply that it would be beneficial to source probiotic therapies or other microbiome treatments based on the geographic origin or genetic ancestry of the host. In contrast, the large range of phylogenetic distances in Fig. 2 suggests that such therapies might be better informed by directly genotyping the host’s microbiota (rather than the host), or by identifying smaller sub-regions of microbial genomes that exhibit stronger geographic associations (25). In the latter case, lower genome-wide levels of codiversification could make it easier to detect such locally adapted regions, by searching for parallel genes or haplotypes across phylogenetically distinct genomic backgrounds (20, 26, 27). Stronger signals of codiversification might also be present in human populations with particularly strong and/or ancient geographic separation (5, 8), which could reduce the rates of microbial gene flow at the largest geographic scales. Further efforts to explore these possibilities represent promising avenues for future work.

In light of the above results, it is also interesting to ask why the bacterial phenotypes that were most associated with codiversification in Ref. (9) (e.g. genome size or poor growth at 27^*°*^C) have not led to stronger genetic differences in certain species, given their dramatically reduced ability to survive outside their hosts. While species with the strongest phylogenetic correlations were enriched in both of these phenotypes, other prominent commensals with smaller genomes (e.g. *Bifidobacterium longum*; Fig. 2D, top) or similar temperature sensitivities (e.g. *Parabacteroides merdae*; Fig. 2D, bottom) exhibited some of the lowest correlations with their host genomes. These contrasting examples suggest that subtle differences in the rates of gene flow and recombination could be a major driver of these large-scale geographic patterns, even if they are not directly related to the historical process of codiversification. Understanding the mechanisms that allow commensal gut bacteria and spread between hosts – and the signatures this leaves in their genomes – will be critical for furthering our understanding of how humans and their microbiota have coevolved.

## Acknowledgments

I thank T. Suzuki, N. Garud, J. Sonnenburg, J. Lopez, Z. Liu and other members of the Good lab for useful discussions. This work was supported in part by the Alfred P. Sloan Foundation grant FG-2021-15708 and NIH NIGMS Grant No. R35GM146949. B.H.G. is a Biohub, San Francisco, Investigator.

## Data and materials availability

Phylogenies and alignments from Ref. (9) were downloaded from the Dryad repository provided in that work; host and bacterial metadata were obtained from the Supplementary Materials. All analysis code used in this work is available on Github (https://github.com/bgoodlab/codiversification_microbiome).

## Supplemental Methods

### Source Data

Phylogenetic trees and alignments from Ref. (9) were downloaded from the associated Dryad repository (28). Host and bacterial metadata were obtained from the Supplementary Materials of Ref. (9). Manual examination of these data revealed the host genomes of three Cameroonian subjects were highly diverged from other Cameroonians and from the rest of the study cohort (Fig. S2). Since the genomes of their gut bacteria were not similarly diverged, these individuals were removed from all analyses except Fig. S2B so that they would not inflate the phylogenetic correlations between different halves of the human genome.

### Phylogenetic correlations within the human genome

Comparisons between different halves of the human genome were performed by splitting the human genotype alignments into two equal subsets. Phylogenies were re-inferred for each set using SNPhylo (29), following the same procedures and parameters described in Ref. (9). Phylogenetic distances were extracted form the corresponding phylogenies, while raw genetic divergences were computed as the fraction of mismatches within the subset of alignable sites for each pair (Fig. S4). PACo scores were computed using a modified version of the code provided by Ref. (9).

### Phylogenetic correlations in simple models of geographic structure

To demonstrate that phylogetic correlations can arise even in the absence of codiversification, we considered two hypothetical models of non-codiversifying bacteria that are illustrated in Fig. 1.

In the first model, bacterial genomes were sampled from the phylogeny in Fig. 1A: genomes from the same clade had a star-like phylogeny, with an internal phylogenetic distance of *d*_*i*_ ≈ 0.3, while branch lengths separating the two clades were set to *d*_*o*_ ≈ 1. For each of the 521 subjects in the original dataset that possessed a *Prevotella copri* strain, we randomly sampled a hypothetical microbial genome from one of the two clades in the relative proportions illustrated in Fig. 1A (*p*_*α*_ ≈ 0.7, 0.6, and 0.3, for Africa, Europe, and Asia respectively). These proportions were purposefully chosen to have a different topology than the genetic relatedness between the three human populations, to emphasize that they are independent of human ancestry. This hypothetical microbial phylogeny was then combined with the observed human phylogeny to calculate an equivalent PACo score, using the same methods described above.

In the second model, we simulated the opposite extreme of a rapidly recombining population with finite rates of geographic mixing. We considered a standard neutral island model with three equal-sized populations representing the three different continents in Fig. 1. Each continent was assumed to be perfectly well-mixed, while migration between continents occurred at a uniform per capita rate *m*. Recombination was assumed to occur sufficiently rapidly within each continent that the mutations at each locus could be sampled independently. This allowed us to simulate the model in a more efficient manner by performing a large number of single-locus simulations across a range of population-scaled migration rates (*Nm*). We assumed an infinite sites limit, in which each sampled polymorphism was founded by a single mutation event. For each value of *Nm*, a sample of hypothetical microbial genomes was constructed by sampling *L* = 2 *×* 10^4^ independent alleles from the simulated matrix of allele frequencies across continents. The number of polymorphisms (*L*) was chosen to coincide with the typical genetic diversity of human gut bacteria (19).

From these simulated microbial genome alignments, we inferred a genome-wide phylogeny using RAxML (30) using the same procedures described in Ref. (9). We then combined this simulated phylogeny with the observed human phylogeny to estimate PACo scores in Fig. 1B. We also performed a standard Mantel test (31) using the matrix of phylogenic distances. For both metrics, we observed statistically significant phylogenetic correlations with the human genome even under high scaled migration rates (*Nm* ≫ 1). In this limit, most sampled genomes will have recently migrated from another continent, on a timescale much shorter than the typical TMRCA (∝ *N*).

### Estimates of the bacterial molecular clock

To estimate the bacterial genetic divergence accumulated since the out-of-Africa expansion, we used two different estimates of the bacterial molecular clock. Direct observations in *E. coli* (13) and *B. fragilis* (14) are consistent with *in vivo* substitution rates of *r* ∼ 10^−7^ − 10^−6^ per bp per year. Compounded over ∼60,000 years, this leads to an estimated divergence of

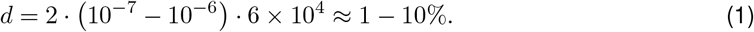

at neutral loci (e.g. synonymous sites) between isolated pairs of strains. Similar (though slightly cruder) bounds can also be estimated from first principles using mutation rates measured in *in vitro* mutation accumulation experiments (∼10^−9^ − 10^−10^ / bp / gen (32)). Given an estimated growth rate of 1-10 generations per day (33), this leads to a an estimated substitution rate of *r* ∼10^−7.5^ − 10^−5.5^ per bp per year, which increases the range of *d* by a factor of 3 in each direction.

All analysis code used for data processing and figure generation is available on Github (https://github.com/bgoodlab/codiversification_microbiome).

## Supplemental Figures

**Figure S1:**
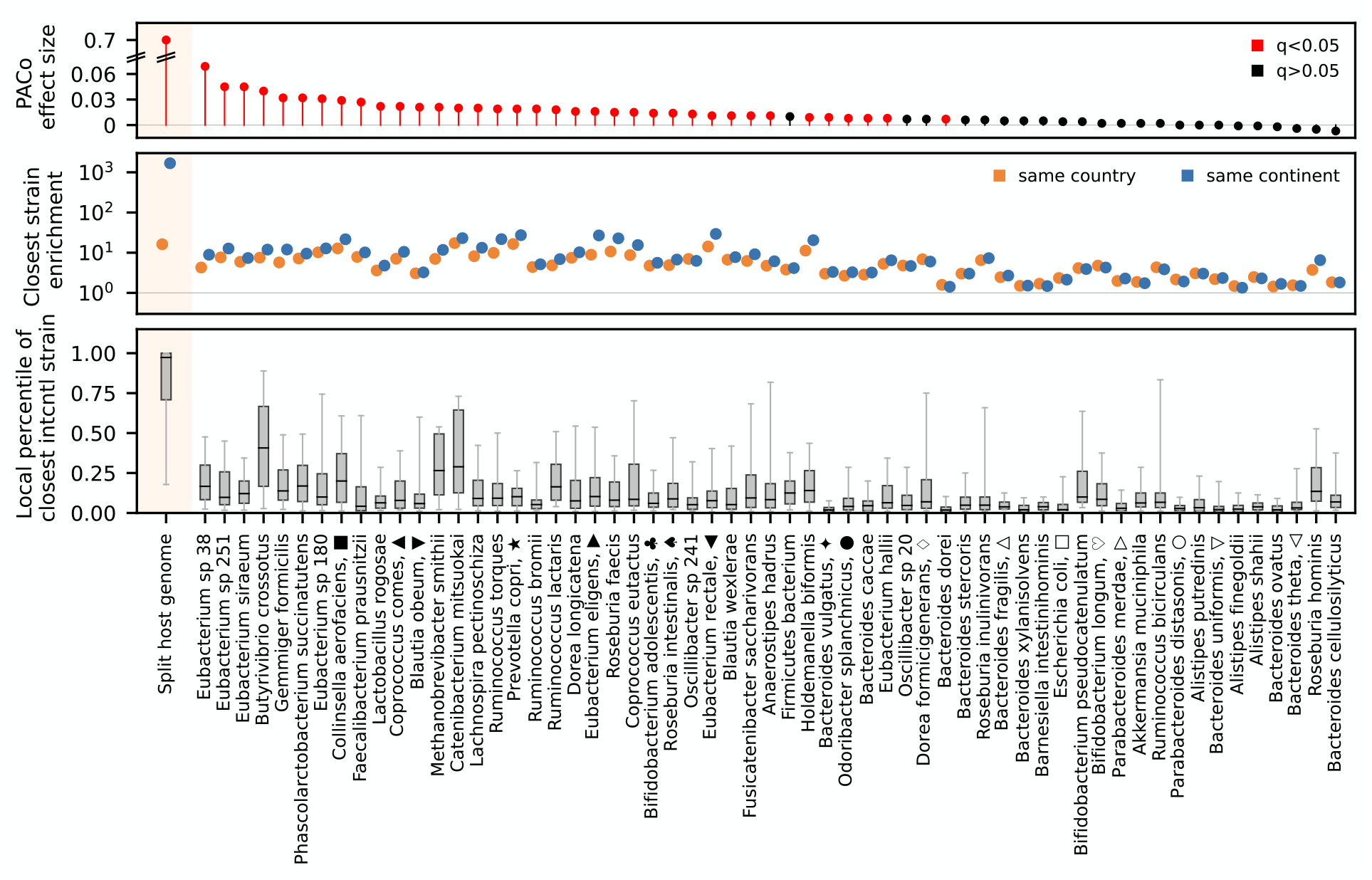
Closely related bacterial strains within and across continents. Top: PACo effect sizes for all of the species analyzed by Ref. (9); species with significant q-values are highlighted in red. Note the large discontinuity between the human and bacterial effect sizes (hatches). Middle: Enrichment of the probability that the closest relative of a strain derives from the same country (orange) or continent (blue). The enrichment factor is calculated as *e* = *f* (1 − *f*_0_)*/*(1 − *f*)*f*_0_, where *f* is observed fraction of strains whose closest relative is located in the same country (or continent) and *f*_0_ is the expected fraction if the strains were randomly permuted across countries. These data show that closest bacterial strains are more likely to come from the same country or continent, but less frequently than their human hosts. Bottom: the phylogenetic distance percentile of the closest inter-continental strain, relative to the set of strains from the same country. These data show that most strains have close relatives on other continents (e.g. in the top quartile of the within-country distribution).

**Figure S2:**
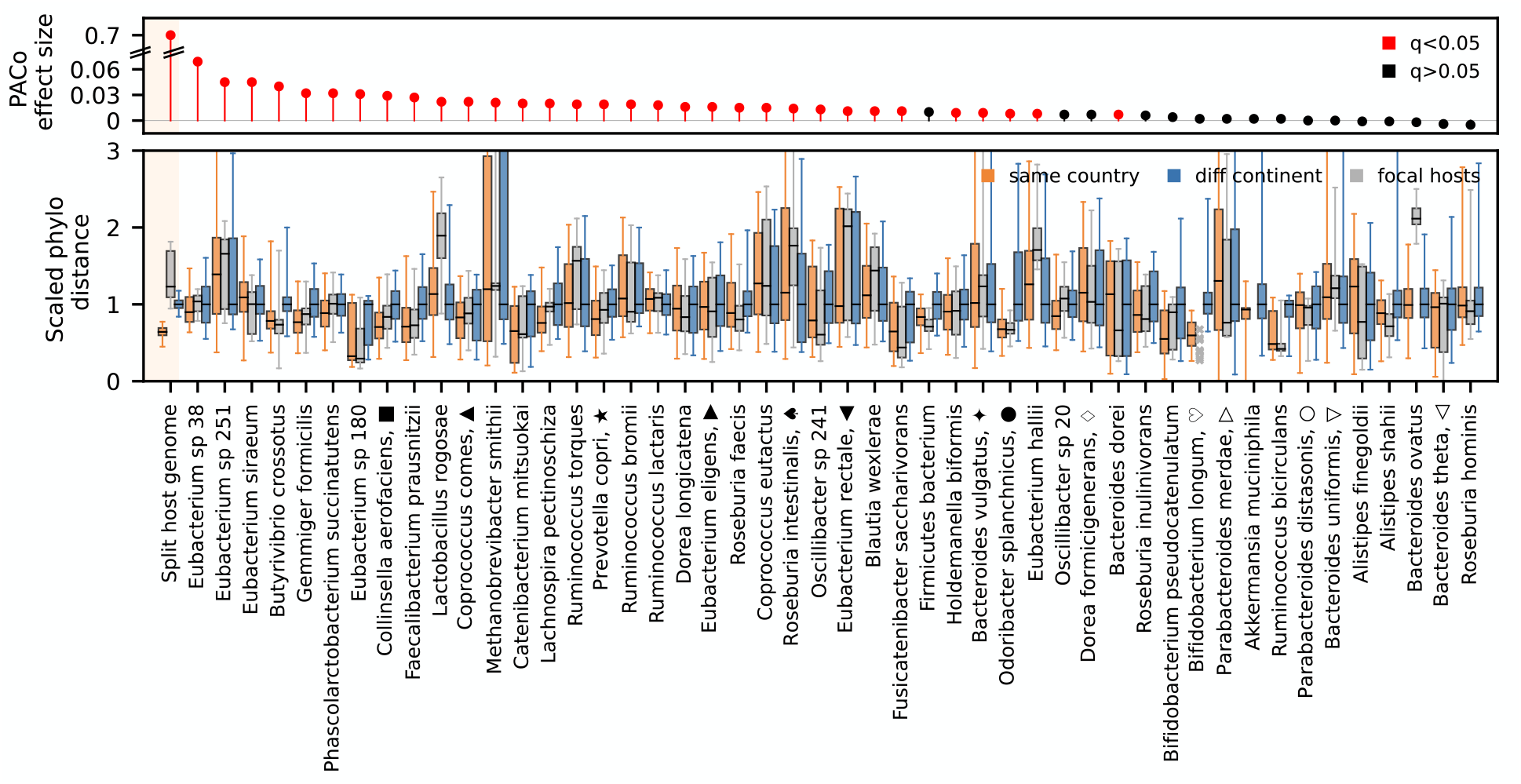
Limited signatures of codiversification in the most highly diverged hosts. Three hosts from Cameroon (BEB30, BEN18, BEN43) were highly diverged from other Cameroonians and from the rest of the human population (bottom, left). The bottom panel shows that for most species, the divergence of their gut bacteria was comparable to the local strain diversity within Cameroon, regardless of the species’ PACo effect size (top). To aid visualization, the phylogenetic distances in the bottom panel are scaled by the median inter-continental distance within each species.

**Figure S3:**
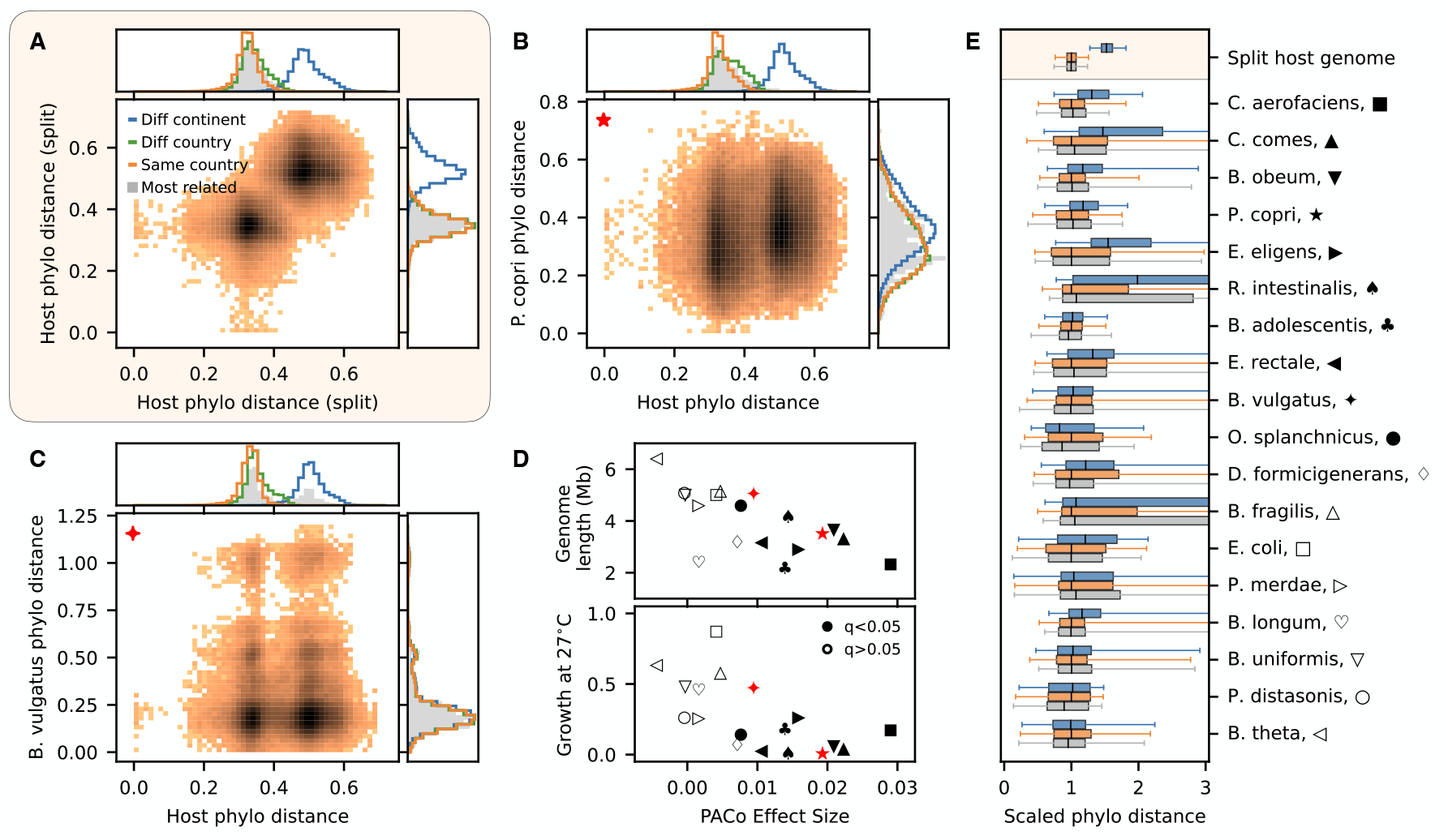
Analogous version of Fig. 2 computed after randomizing host labels within continents.

**Figure S4:**
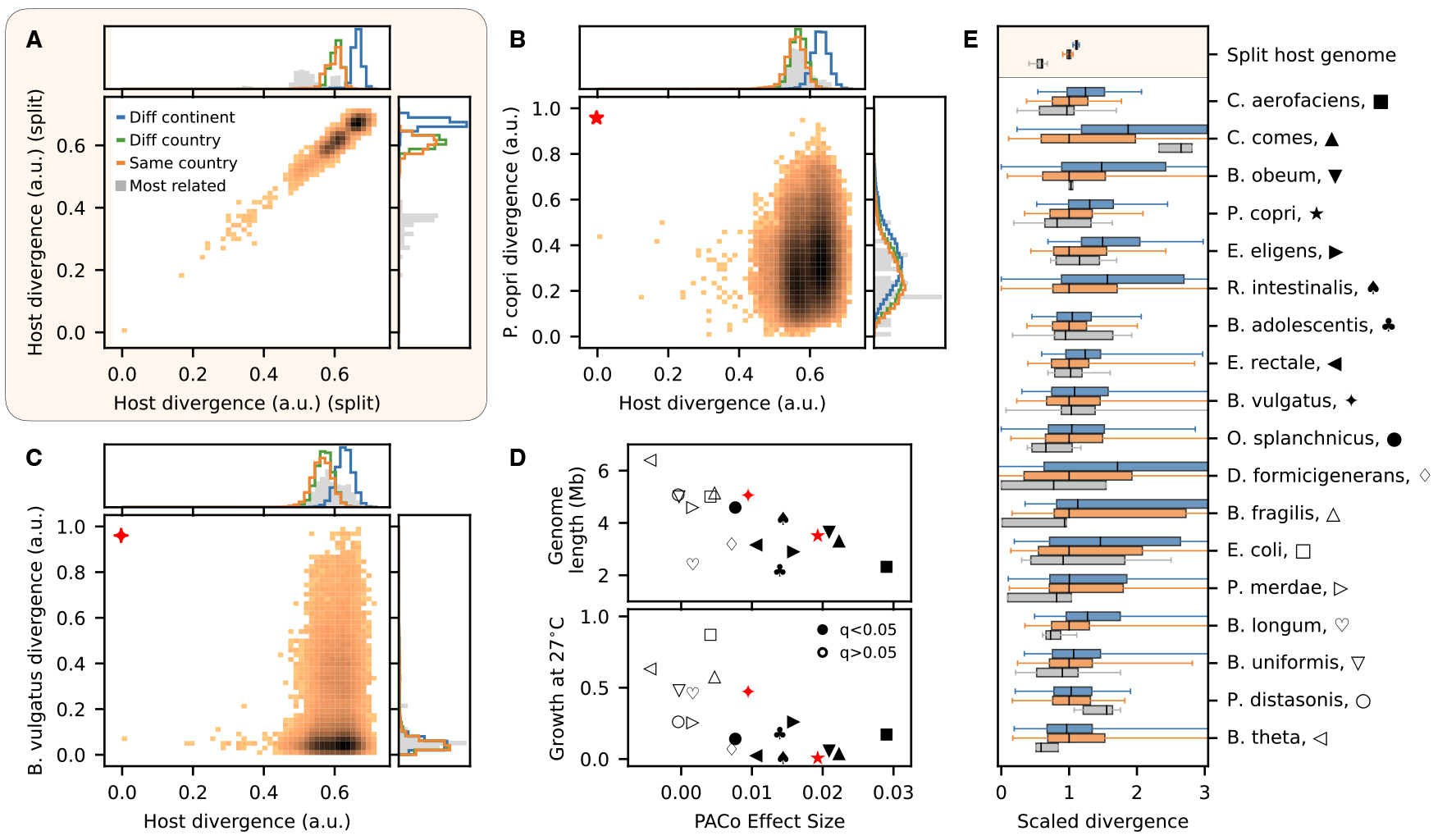
Analogous version of Fig. 2 computed using raw genetic divergences rather than inferred phylogenetic distances.

